# Suzetrigine-evoked analgesia is potentiated by Nav1.7 null opioid-related mechanisms

**DOI:** 10.64898/2026.10.05.755985

**Authors:** Eva Villalba Riquelme, Olena Didenko, Sam Gossage, Jorg Isensee, Tim Hucho, John N Wood

**Affiliations:** Molecular Nociception Group, Wolfson Institute for Biomedical Research, UCL, Gower Street London WC1E 6BT, UK; Translational Pain Research, Department of Anaesthesiology and Critical Care Medicine, Faculty of Medicine and University Hospital Cologne, University Cologne.

## Abstract

The genetic evidence for an essential role of sodium channel Nav1.7 in human pain conditions is compelling. Unfortunately, potent selective channel blockers of Nav1.7 are not useful analgesics because they have deleterious side effects, particularly on the autonomic nervous system. Here we have evaluated the downstream mechanisms induced by Nav1.7 loss *in utero*. Nav1.1 compensates for loss of Nav1.7 in sensory neurons and antisense Nav1.1 oligonucleotides diminish excitability in Nav1.7 embryonic null sensory neurons. Enhanced endogenous opioid signalling contributes substantially to analgesia. This is linked to altered gene expression that potentiates opioid signalling through lowered levels of proteins that terminate G-protein signalling (RGS4 and RGS8). There is also enhanced opioid drive through increased enkephalin levels linked to lowered sodium levels in Nav1.7 null neurons. Opioid signalling is the main driver of analgesia, but enhanced potassium channel expression (KCNC4 KCNK2, KCNMB2, KCNAB3) all implicated in pain control, may also contribute to the small level of naloxone-resistant analgesia. These findings suggest a potential new route to analgesia, through mimicking the events at Nav1.7 null sensory neuron terminals. Here we demonstrate potentiation of suzetrigine-evoked analgesia with RGS antagonists alone or in combination with normally inactive doses of morphine in formalin-evoked pain and CFA-induced inflammatory pain. Cancer pain is under investigation. These data are of potential clinical interest.

## Introduction

Progress in treating pain has been slow. Extrapolation of drug studies on inbred rodents to outbred humans have often failed, even though all human analgesics work well in mice. It is clear that peripheral drive from nociceptors is key to most pain conditions ^1^. Drugs that block pain exert their actions on these cells in terms of blocking action potential propagation (lidocaine) ^2^ , blocking the hyperalgesic effects of inflammatory mediators (NSAIDs) and blocking neurotransmitter release (opioids and gabapentinoids)^3,4^. The major problem with present analgesic drugs seems to be undesirable side effects rather than lack of efficacy, as demonstrated most dramatically by the recent US opioid epidemic ^4^ that has cost hundreds of thousands of lives.

Recently a major advance was made with the development and FDA approval of a selective allosteric Nav1.8 sodium channel blocker, Suzetrigine^5^. Blocking nociceptor input is potentially effective , but other sodium channels present in Nav1.8+ neurons can contribute to nociceptive drive and some conditions, for example human cancer-induced bone pain, are Suzetrigine-resistant ^6^. Although Suzetrigine is remarkably specific for human Nav1.8, higher doses used here also inhibit the mouse channel ^7^. The binding site defined by mutagenesis and domain-swaps is in the region of human sequence KKGS found in the extracellular S3-S4 loop domain 2. The same region in mouse is homologous but adjacent residues marked in red differ (LEL<u>STS</u>KKGSLSVLR<u>T</u>FRLL – Mouse, LELGVAKKGSLSVLRSFRLL – Human). These changes, potentially contributing to subtly different conformations, may explain the 1000-fold lower affinity for the rodent channel ^5,7^.

Another sodium channel that is important for human pain is Nav1.7. Mice and humans that lose expression of functional Nav1.7 *in utero* develop into pain-free adults^8,9^. This led to a great effort to produce selective Nav1.7 antagonists that would mimic the loss-of-function genetic mutants. Unfortunately, Nav1.7 is very broadly expressed in the central nervous system and sensory and autonomic neurons. Hence selective Nav1.7 channel blockers have life-threatening side effects that preclude their development for human use ^10^, despite their potent analgesic actions when delivered to central terminals of sensory neurons ^11^. This raises the interesting question of why loss-of-function mutants remain viable and apparently normal, apart from loss of pain and olfaction^12^. A possible explanation is that opioid signalling is a key element in Nav1.7 null analgesia^9^. Opioid receptors are known to be restricted to nociceptive sensory neurons but are not present in autonomic neurons^13^ (see supplementary Table 1)..

In this study, we first explored the mechanisms that rescue the electrical excitability of null Nav1.7 mutants. We examined altered RNA and protein expression in Nav1.7 null peripheral neurons in order to define potential compensatory mechanisms that could rescue the deleterious aspects of the Nav1.7 null phenotype. We have found and validated a mechanism for the rescue of neuronal excitability by Nav1.1 in peripheral neurons from Nav1.7 null embryonic mutants.

Next we explored aspects of opioid signalling in these neurons to identify endogenous mechanisms that could be potentially exploited as new routes to effect analgesia, We identified potential mechanisms for enhanced endogenous opioid mediated analgesia, including potentiating effects on G_0 i_nhibition of pain pathways^14^ and potassium channel induction that may limit excitability of sensory neurons ^15^.These insights allowed us to explore Nav1.8 inhibition combined with molecular mechanisms linked to Nav1.7 null analgesia that do not use channel blockers. Here, we present evidence that these downstream aspects of Nav1.7 opioid-signalling can synergise with the analgesic block of Nav1.8 produced by suzetrigine, and provide a potentiated route to analgesia that may be explored clinically.

## RESULTS

### Nav1.1 functional compensation for Nav1.7 loss in null mutant mice

Increased levels of Nav1.1 have been identified in sensory neurons from embryonic null Nav1.7 null mice ^16^. Nav1.7 also plays an important role in sympathetic neuron function and is the site of action of Nav1.7 antagonist channel blockers, and a principal reason for their undesirable side effects ^10^ .

We used antisense oligonucleotides to down regulate Nav1.1 expression in Nav1.7 null neurons (Supplementary Figure 1) and examined the effect on excitability *in vivo* using GCaMP imaging of somatic responses to noxious stimuli (Figure 1). It is important to remember that Nav1.7 null neurons show entirely normal excitability in vivo with GCaMP imaging or using electrophysiological measurements ^9^ and see Figure 1,2. We also studied excitability *in vitro* using electrophysiological measures (Figure 2). It is clear that there is diminished responsiveness to noxious stimuli when Nav1.1 is knocked down by antisense oligonucleotides in Nav1.7 null mutant neurons. In terms of GCaMP imaging of live null mice DRG, we found that sensory neuron somata depolarisation was much reduced in response to heat and mechanical stimulation whilst cold responses were little altered. We summed the GCaMP responses to noxious stimuli in fig 1l and showed a major loss of responsiveness in response to the knockdown of Nav1.1. In addition, the remaining calcium responses to noxious stimuli were much reduced as shown in the violin plots in Figure 1 m,n,o) .

**Figure 1.**
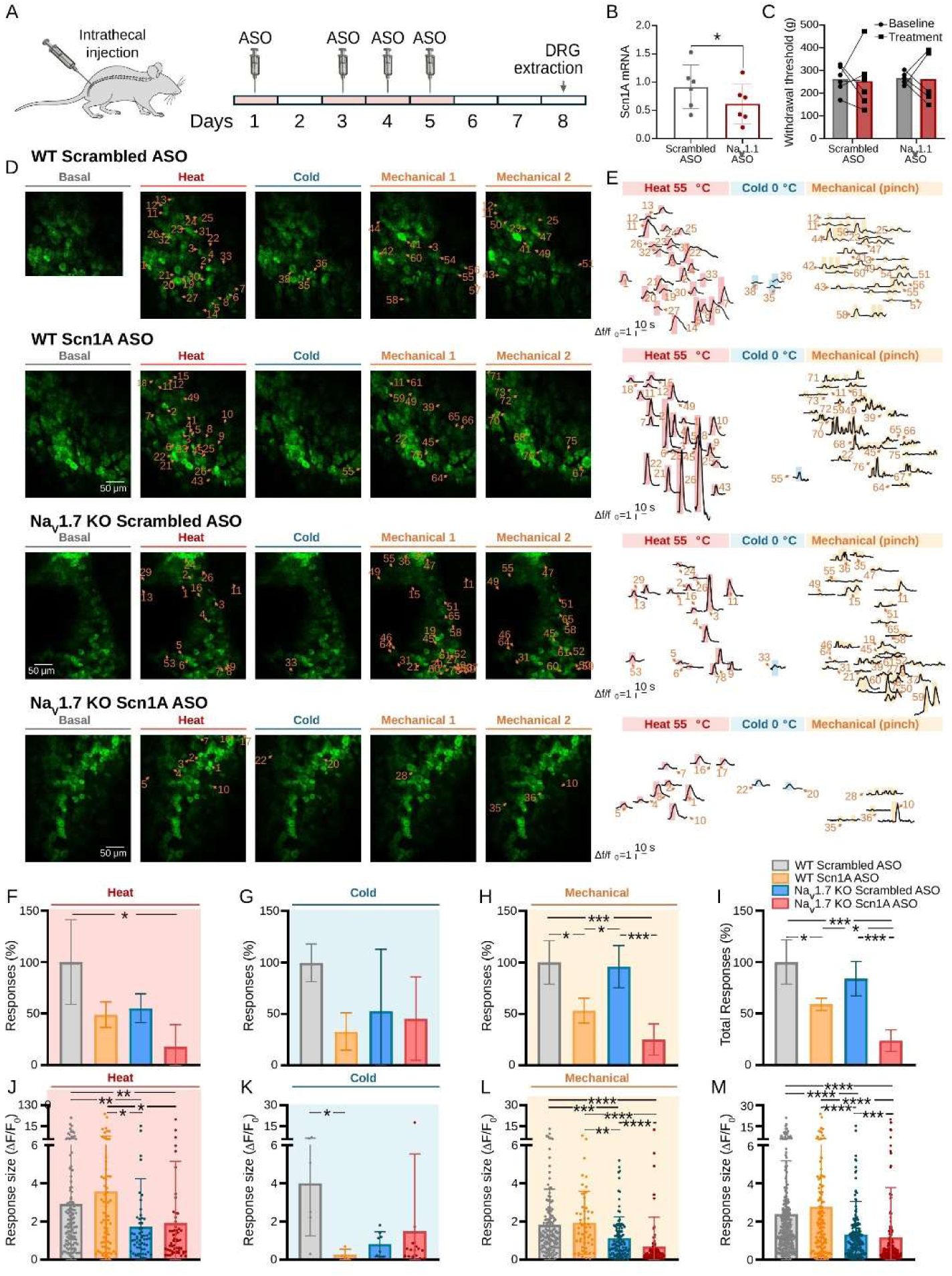
Na_V1_.1 is essential for maintaining *in vivo* noxious calcium responses in embryonic Na_V1_.7 null DRG neurons. **A)** Diagram of the experimental paradigm followed for these experiments. Antisense oligonucleotides to downregulate Na_V1_.1 expression and its scrambled control were injected on days 1, 3, 4 and 5. Experiments were performed 72-96h after the last injection (days 8-9). **B)** Treatment with antisense oligonucleotides significantly reduced Scn1A mRNA levels on lumbar DRG L2-L6. Relative expression normalized to the housekeeping gene Rpl29. Unpaired t-test. **C)** Scn1A downregulation did not affect pain threshold in the Randall-Selitto test. The tail withdrawal threshold was measured at day 1 before injections and day 8 prior to experiments. Unpaired t-test. **D)** Example images showing calcium responses to noxious heat (55°C water), cold (0°C water) and mechanical stimuli (pinch) applied to the ipsilateral paw of the mouse for WT treated with scrambled ASO or with Scn1A ASO or for Na_V1_.7 KO mice treated with scrambled ASO or Scn1A ASO. Responding cells are numbered and indicated with orange arrows. **E)** Traces of the fluorescence changes (ΔF/F_0)_ detected in response to the noxious stimuli for the cells shown in the images indicated on the left panels. **F-I)** Percentage of neurons that responded to noxious heat **(F)**, cold **(G)**, and mechanical **(H)** stimuli or to all the noxious stimuli applied **(I)** normalized by the total number of neurons that responded to noxious stimuli per area in the WT mice treated with scrambled ASO. One-way ANOVA with Tukey’s multiple comparisons test. **J-M)** Scatter dot plots of the calcium response size (ΔF/F_0)_ for noxious heat **(J)**, cold **(K)**, and mechanical **(L)** stimuli, or to all the noxious stimuli applied **(M)**. Kruskal-Wallis with Dunn’s multiple comparisons test. * p ≤ 0.05. **p ≤ 0.01. ***p ≤ 0.001. **** p ≤ 0.0001.

**Figure 2.**
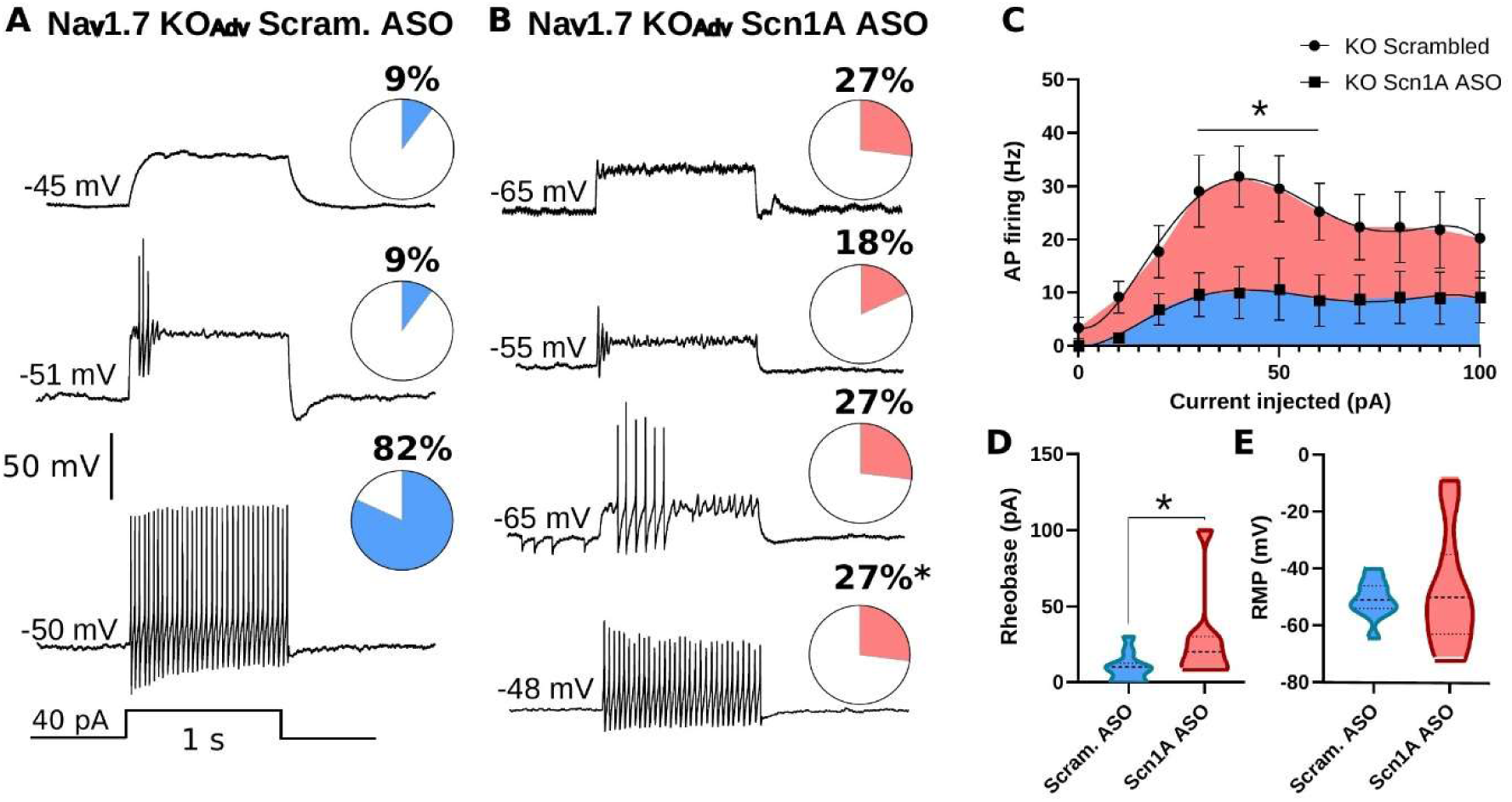
Naᵥ1.1 ASO treatment reduces evoked action potential firing in small Naᵥ1.7 null DRG neurons. (A and B) Representative action potential recordings evoked by a 40 pA, 1 s current injection in neurons from Naᵥ1.7 KOᴬᵈᵛ mice treated with scrambled ASO (A) or Scn1a ASO (B). The proportion of neurons showing each firing pattern is given to the right of each trace. Fisher’s exact test. (C) Action potential firing frequency evoked by current injections from 0 to 100 pA in 10 pA steps. Scn1a ASO treatment reduced firing frequency. Symbols show the mean and bars the SEM. (D) Rheobase, which was increased after Scn1a ASO treatment. Mann–Whitney U test. (E) Resting membrane potential. Lines show mean ± SD. Unpaired t test. *P < 0.05.

With *in vitro* electrophysiological studies from Nav1.7 null mice treated with antisense oligonucleotides to Nav1.1, we found that the percentage of neurons responding strongly to depolarising input fell from 82% to 27% with down regulation of Nav1.1, whilst unresponsive neurons increased from 9 to 27% (Figure 2).

### Sodium and Nav1.7

Nav1.7 is expressed at high levels in sensory neuron terminals, but at low levels in the somata and axons *in vivo* – (see figure 1 in reference ^17^) . We wanted to measure the effect of deleting Nav1.7 on sodium concentrations in the central terminals of mice, but this proved to be technically very challenging. However, examining cultured sensory neurons from control and embryonic null Nav1.7 mice showed that there was a difference in intracellular sodium levels as measured with SBFI (Figure 3a). In all size classes of neuron small <25 micron diameter, medium 25-40 micron diameter an large >40 micron diameter. the intracellular sodium levels were close to 10mM – about half that found in wild type mice (Figure 3).

**Figure 3.**
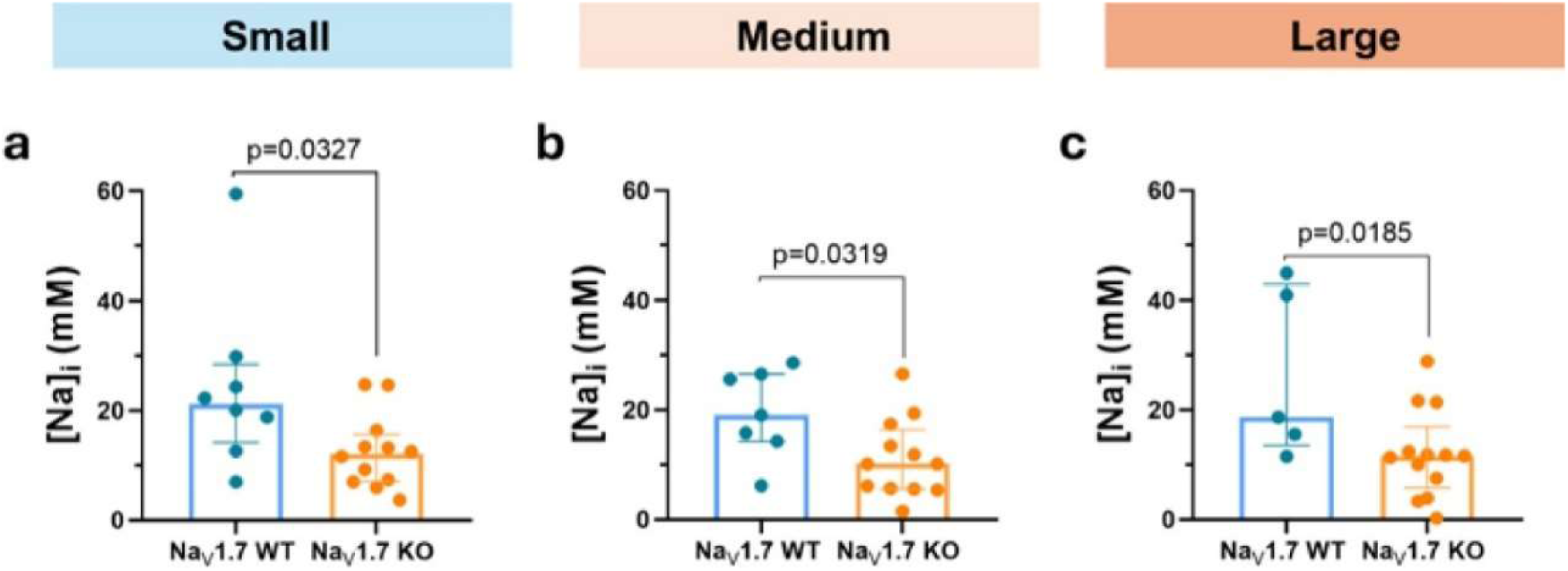
Intracellular sodium is lower in Naᵥ1.7 embryonic null sensory neurons. Intracellular sodium concentration, estimated with the sodium indicator SBFI, in cultured DRG neurons from Naᵥ1.7 wild-type and Naᵥ1.7 embryonic null mice, shown separately for small (a; diameter < 25 µm), medium (b; 25 µm ≤ diameter < 40 µm) and large (c; diameter ≥ 40 µm) neurons. In all three size classes the intracellular sodium concentration in null neurons was close to 10 mM, about half that measured in wild-type neurons. Each point is the mean of all neurons of that size class on one coverslip. Bars show the median and interquartile range. Unpaired t test; P values are given on the panels.

Given the link between intracellular sodium levels and enkephalin production in sensory neurons ^18^, it was of interest to investigate the potentiated opioid receptor signalling in Nav1.7 null sensory neurons demonstrated by ^19^ and ^9^ . Immunoprecipitation of Nav1.7 interacting proteins identified the G-protein Gαo as an interactor^16^. This is significant as the Gαi/o-family of heterotrimeric G proteins are linked to mu-opioid receptors (OPRM1) that mediate analgesia. OPRM1 receptors are known to be inhibited through a sodium binding site. Previously, we have shown that blocking Nav1.7 with high concentrations of tetrodotoxin (TTX) can induce increased PENK mRNA production ^18^. In addition, measuring activity of type II protein kinase A (PKA-II) downstream of G-protein / adenylyl cyclase signalling, revealed potentiated opioid receptor signalling in isolated sensory neurons of Nav1.7 null mutant mice ^20^. Here we used this assay to test for changes in opioid receptor signalling after long term TTX pharmacological block of Nav1.7. In a first experiment, we measured the inhibitory effect of the OPRM1-selective opioid fentanyl on forskolin-induced PKA-II activity after preincubation with high doses of TTX (6 h, 500 nM) (Figure 4b). Preincubation with TTX did not change the inhibitory effect of fentanyl suggesting that altered intracellular levels of sodium do not necessarily enhance opioid signalling . The TTX dose was effective as veratridine-induced PKA-II responses were suppressed by TTX with an IC50 of 60 nM ^21^. This aligns with a second experiment (figure 4a) in which we observed no effect of TTX (300 nM, 24 h) or PF-04856264 (1 μM, 24 h) on the inhibitory action of the endogenous opioid peptide Met-enkephalin (Figure 4 c, d) on forskolin- or serotonin-induced PKA-II activity ^20^.

**Figure 4.**
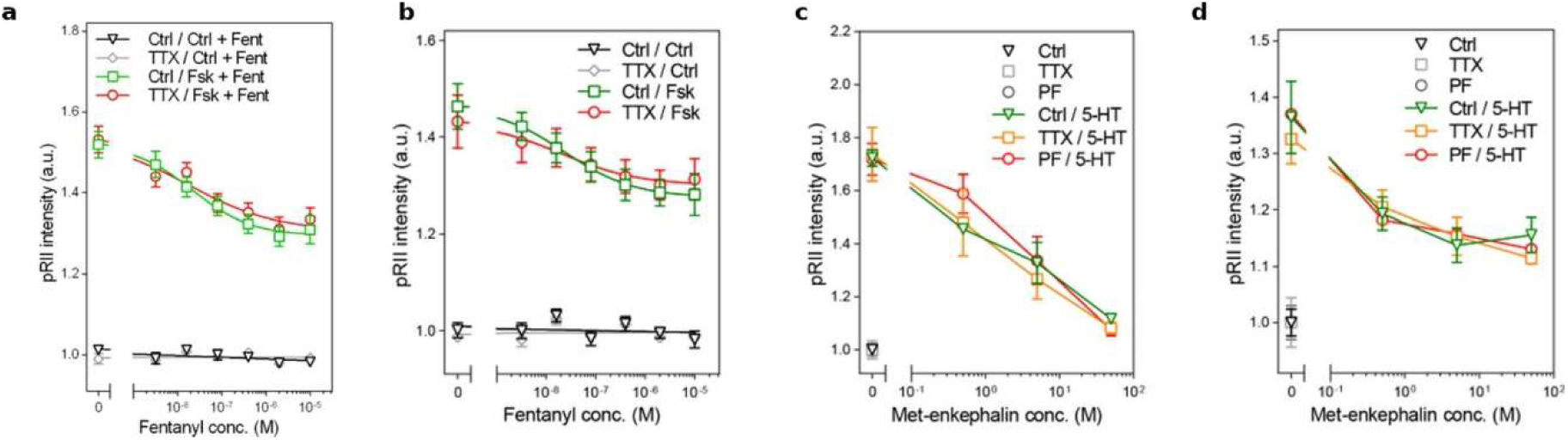
Tetrodotoxin block of Naᵥ1.7 does not potentiate opioid receptor signalling in sensory neurons. (a and b) Inhibitory effect of the OPRM1-selective opioid fentanyl (0 to 10 µM) on forskolin-induced (Fsk; 3 µM, 3 min) PKA-II activity after preincubation with tetrodotoxin (TTX; 500 nM) for 6 h. In (a) fentanyl was added to all conditions; (b) shows the same preincubation and stimulus conditions. Preincubation with TTX did not change the inhibitory effect of fentanyl. DRG neurons are from n = 8 mice (4 female, 4 male), analysed across two independent experimental days. (c and d) Inhibitory effect of the endogenous opioid peptide Met-enkephalin (0, 0.5, 5 and 50 µM) on forskolin-(3 µM, 3 min) or 5-HT-induced PKA-II activity after preincubation with TTX (300 nM, 24 h) or PF-04856264 (PF; 1 µM, 24 h). DRG neurons are from n = 4 mice (2 female, 2 male), analysed across two independent experimental days. Symbols show the mean and error bars the SEM. pRII, phosphorylated RII subunit of PKA-II.

### Pain-related gene expression in Nav1.7 embryonic null sensory neurons

As altered intracellular levels of sodium do not appear to contribute to enhanced opioid signalling, this event must rely on another mechanism. We therefore examined changes in gene expression in the Nav1.7 null sensory neurons using both microarray and mass spectrometry data (see Table 1). We focused in particular on proteins known to be involved in regulating opioid signaling or sensory neuron excitability. One regulatory step that is key to opioid action is the control of G-protein signalling. GPCR signalling is deactivated by intrinsic GTPase activity present in the α–subunit that terminates downstream signalling pathways. The GTPase activity itself is regulated by other proteins, in particular the class of regulators of G-protein receptor signalling (RGS) proteins play an important role ^22^. Pain signalling is known to be regulated by these proteins ^22^ . For example, genetic deletion of RGS4 has profound effects on pain signalling in rodents, eliminating some aspects of second phase formalin-induced behaviour and reversing cold and mechanical hypersensitivity induced by inflammatory and neuropathic stimuli ^23^ . Perhaps more importantly, pharmacological block of RGS action potentiated opioid signalling a 100-fold^20^. We found that RGS4 and RGS8 were indeed downregulated in sensory neurons in Nav1.7 null mice ^24^. It was thus plausible that the lowered levels of RGS proteins present in Nav1.7 null mice could contribute to enhanced opioid receptor signalling. The use of a RGS4 inhibitor is consistent with this suggestion. The Kim group showed that the covalent RGS4 inhibitor CCG50014 massively potentiated opioid analgesic effects ^25^, as well as reducing a variety of nociceptive responses.

**Table 1.**
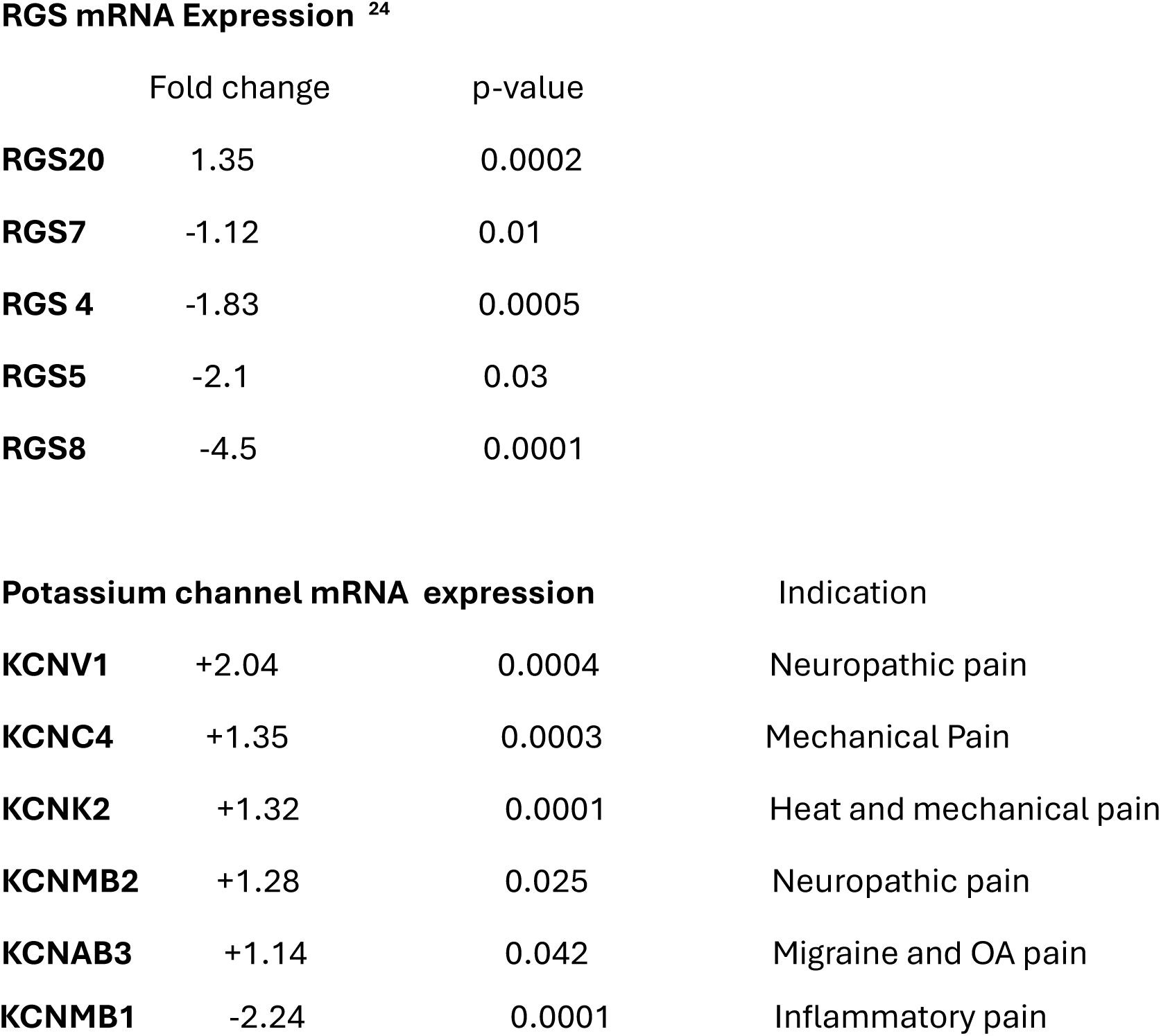
Pain gene candidates regulated in Nav1.7 null sensory neurons.

Some elements of Nav1.7 null analgesia are not reversed by the opioid antagonist naloxone. We found upregulation of some potassium channels linked to pain that may also contribute to regulation of excitability (Table 1)

These observations suggested that the useful analgesia produced by Nav1.8 antagonists might be potentiated with mechanisms associated with embryonic Nav1. 7 null analgesia ^21^. We therefore tested mice in a variety of pain assays using high dose suzetrigine (10 mg/kg) known to be effective in rodents as the Nav1. 8 antagonist alone ^7^, or in combination with the RGS inhibitor CCG50014 ^20,21^ (100 nmol). We also incorporated a low dose of morphine (7.5ug) into the protocol that has no analgesic effects alone at the dose used (^26,27^ ^28^,) as confirmed in Figure 5.

**Figure 5.**
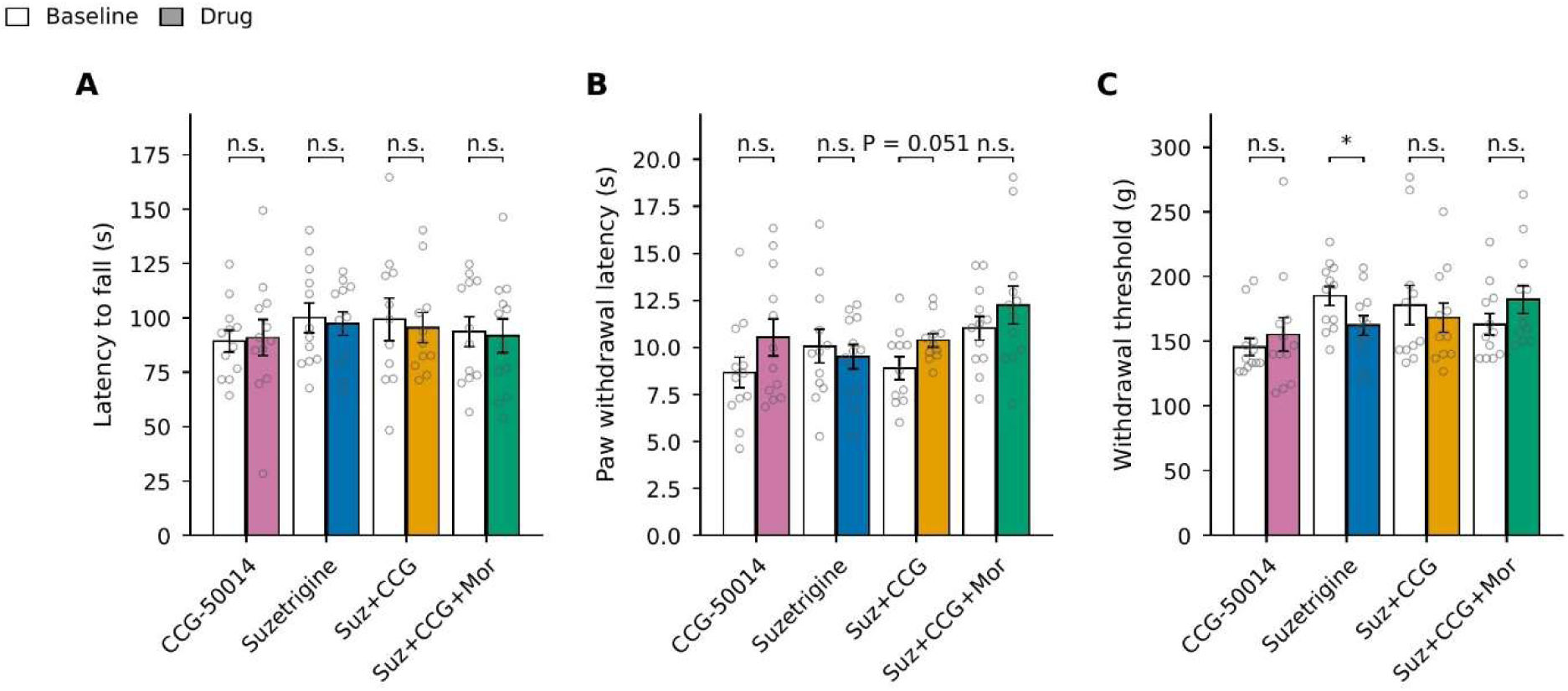
Neither CCG-50014 nor Suzetrigine, nor low dose morphine alter nociceptive thresholds or motor performance in uninjured mice. (A) Rotarod latency to fall. (B) Hargreaves paw withdrawal latency. (C) Randall–Selitto withdrawal threshold, measured on the tail. All measures were taken at baseline (open bars) and after drug administration (coloured bars) in the same animals (47 mice in total; sexes balanced). Bars show the mean, error bars show the SEM and points show individual animals. Exact P values are given where 0.05 ≤ P < 0.08; n.s., not significant; *P < 0.05 (paired two-tailed t test).

**Figure 6.**
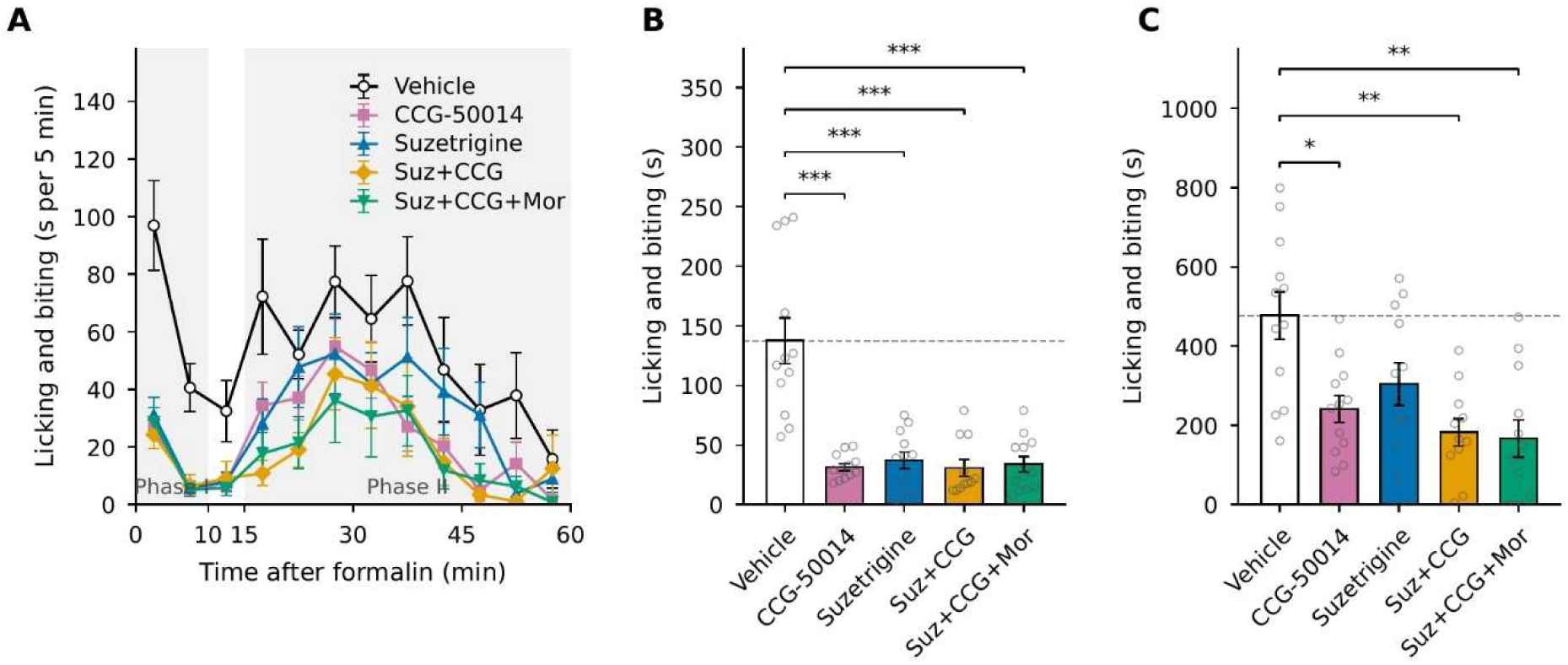
CCG-50014 and suzetrigine reduce formalin-evoked nocifensive behaviour, and their co-administration reduces it further. (A) Time course of licking and biting of the injected paw in 5-min bins. Symbols show the mean and error bars the SEM. Shading marks phase I (0 to 10 min) and phase II (15 to 60 min); the interphase (10 to 15 min) was excluded from both phase totals. (B) Phase I and (C) phase II totals by treatment. Bars show the mean, error bars show the SEM and points show individual animals. The dashed line marks the vehicle mean. n = 12 per group except suzetrigine + CCG-50014 (n = 11); both sexes. *P < 0.05; **P < 0.01; ***P < 0.001 versus vehicle (two-tailed Mann–Whitney U test, Holm-adjusted across the four comparisons within each phase).

Surprisingly, first experiments measuring acute pain thresholds showed that suzetrigine was relatively ineffective (Figure 5) . Similar results have been obtained in rodents by the Zamponi group ^7^. However, in a variety of pain-induced models, analgesia was observed, most clearly in male mice (supplementary Figure 2). This may suggest that activity of mouse Nav1.8 or trafficking enhance suzetrigine binding. The suzetrigine-evoked analgesia was potentiated by blockade of RGS activity with intrathecal CCG50014 suggesting that endogenous opioid signalling could be recruited to further enhance analgesia. With the addition of low-level normally inactive morphine to potentiate endogenous opioid signalling, stronger analgesia was induced. We used a dose of morphine that has been shown to be sub-analgesic ^21^, an action that we confirmed with tests on acute pain (Figure 5).

In the formalin test, we found that both suzetrigine and CCG-50014 were effective in reducing first phase behaviour, whilst second phase behaviour was reduced by both drugs that showed a larger effect when co-administered. The addition of morphine had little further analgesic activity in this test.

Inflammatory pain is easily measured with injections of complete Freund adjuvant (CFA). No drug treatment altered the swelling of the paw after injection (Fig 7A) but enhanced thresholds for heat or mechanical pain were reduced with suzetrigine alone that showed more activity with CCG-50014 and better analgesia with the addition of low dose morphine (Fig 7, D.E). Von Frey thresholds including innocuous mechanosensation showed effects of suzetrigine that were neither enhanced with CCG50014 nor morphine (Fig 7b). This observation can be explained by the fact that most innocuous mechanosensitive neurons do not express opioid receptors so neither endogenous nor exogenous opioids would have an effect.

**Figure 7.**
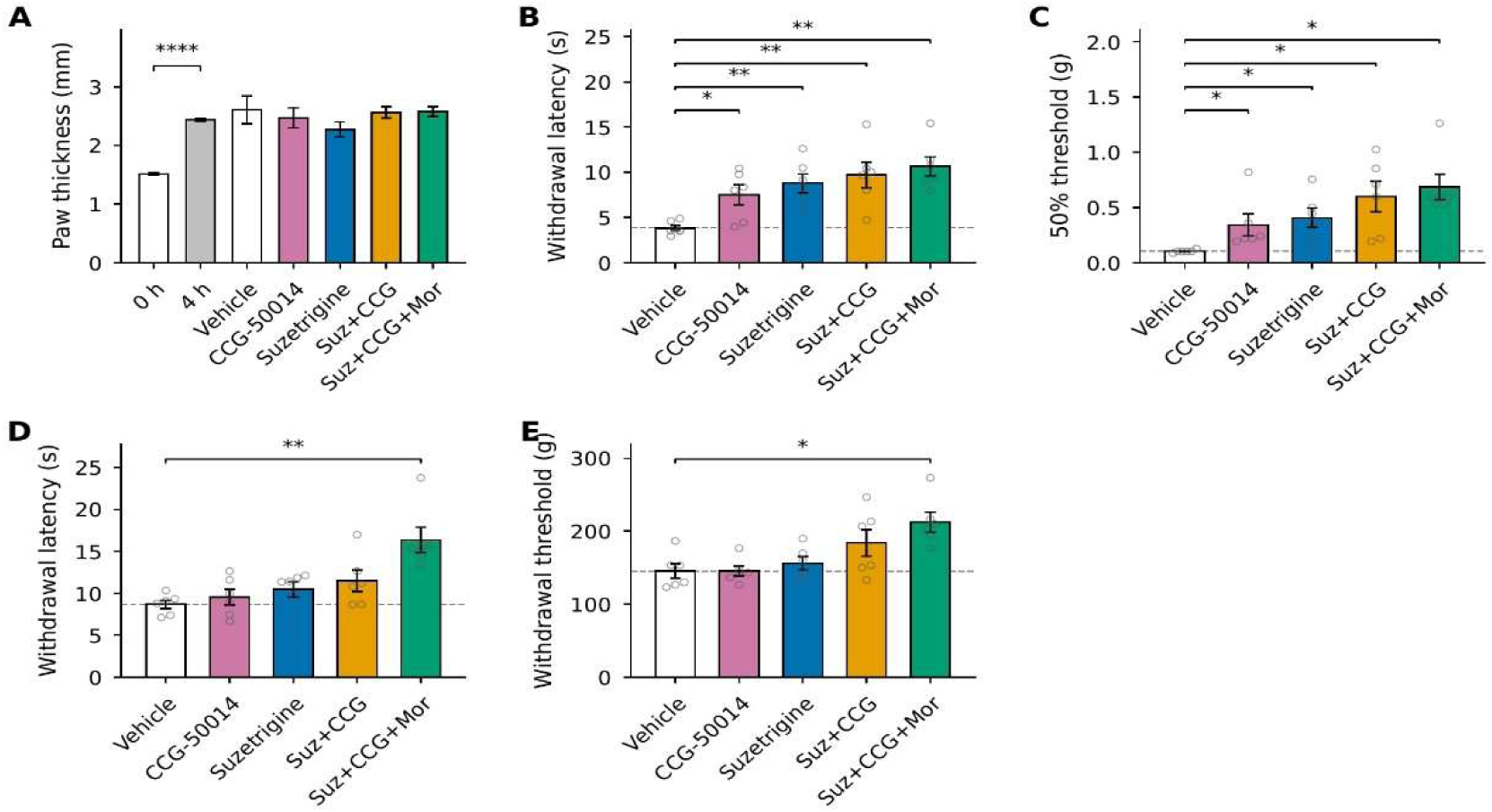
Treatment effects in CFA-inflamed and uninflamed tissue. (A) Paw thickness before CFA (0 h) and at 4 h in the whole cohort, and at 72 h by treatment group. (B) Hargreaves paw withdrawal latency and (C) von Frey 50% withdrawal threshold of the CFA-injected paw at 72 h. (D) Hargreaves paw withdrawal latency of the contralateral, uninflamed paw. (E) Randall–Selitto withdrawal threshold measured on the tail, an uninflamed site. n = 6 per group (male mice). Bars show the mean, error bars show the SEM and points show individual animals. The dashed line marks the vehicle mean. *P < 0.05; **P < 0.01 (B to E, versus vehicle, two-tailed Mann–Whitney U test, Holm-adjusted across the four comparisons within each assay). In (A), ****P < 0.0001 for 0 h versus 4 h (paired two-tailed t test, n = 30).

## Discussion

In this study we have addressed three questions. Firstly, how is function rescued in Nav1.7 null neurons of mice to generate normal excitability? Secondly, what are the mechanisms that lead to endogenous opioid-mediated analgesia in Nav1.7 null mice? Finally, can we potentiate the effects of blocking Nav1.8 activity by suzetrigine with endogenous or exogenous opioid signalling?

The critical role of peripheral sodium channels in nociceptive signalling and pain induction is well established ^17^. In addition, the utility of sodium channel blockers as analgesics is clinically recognised, and lidocaine, a non-specific sodium channel blocker, is well known to abolish pain in dentistry. Nav1.8 shows the remarkable property of selective expression principally in nociceptive sensory neurons.

Examination of 1000 human cadavers has found no evidence for central nervous system expression of Nav1.8 ^5^. Hence specific antagonists of Nav1.8 are likely to have a low probability of side effects. Global deletion of Nav1.8 in mice results in normal fertile mice that have lost elements of inflammatory pain and mechanical pain, further supporting Nav1.8 as a useful analgesic target ^29^

Suzetrigine is an allosteric blocker of Nav1.8 with 30,000 fold selectivity over other sodium channels ^5^. This tour-de-force of medicinal chemistry combined with oral availability has focussed attention on Nav1.8 as a target for non-opioid treatment of human pain. Suzetrigine and the related Latigo-001 compound are both effective in acute pain in humans, and suggestions of utility in diabetic neuropathy pain have been made^30^. The major numerical issue in the pain field is chronic pain associated with old age, particularly associated with mechanical stress in arthritis ^31^.

In this study we have provided evidence that Nav1.1 can rescue excitability in Nav1.7 null neurons. Studies of embryonic null Nav1.7 analgesia where the target is completely deleted have also shown a role for naloxone-repressed endogenous opioid signalling in the resulting analgesia ^9^. The precise mechanisms for Nav1.7 regulation of gene expression remains little understood, although persistent sodium current may play a role ^32,33^ . Enkephalin induction is insufficient to explain this mechanism as NFAT5 deletion leads to enhanced enkephalin production without analgesia ^18^, but potentiated opioid signalling combined with enhanced endogenous enkephalin expression provides a convincing mechanism ^20^. Partial deletion of Nav1.7 in adult animals also causes some analgesia lined to diminished electrical excitability. This study confirmed the fact that enkephalin alone is not sufficient to cause analgesia but ignores opioid receptor signalling potentiation ^5^. We have found that Nav1.1 can replace Nav1,.7 in embryonic null DRG neurons explaining their normal levels of excitability ^10^. The mechanism behind this compensatory channel expression remains unknown. Nav1.7 and Nav1.1 are closely linked on chromosome 2 in both mice and humans. It is known that sodium channels can form dimers ^34^. The lack of Nav1.7 may thus result in more Nav1.1 homodimers. Mass spectrometry has shown that some proteins that interact with Nav1.7 are upregulated in Nav1.7 null neurons ^16^.

The evidence that endogenous opioid signalling can result in useful analgesia is of great interest. Although the Sackler scandal ^35^ of hundreds of thousands of opioid-induced deaths has led to a justifiable reaction against opioid over-prescription, the fact is that exogenous opioids are wonderful analgesics and if used cautiously, they can be of great benefit ^36^. Here we have shown that doses of morphine way below the normal effective dose for analgesia can be useful in potentiating the activity of suzetrigine, an already useful analgesic drug. Perhaps surprisingly, blockade of RGS proteins alone is also effective in causing analgesia. The inhibition of RGS proteins alone must recruit endogenous opioid activity and potentiate suzetrigine activity. These mechanisms are summarised in Figure 8. The inhibitory drug CCG50014, unlike suzetrigine requires intrathecal injection.

**Figure 8.**
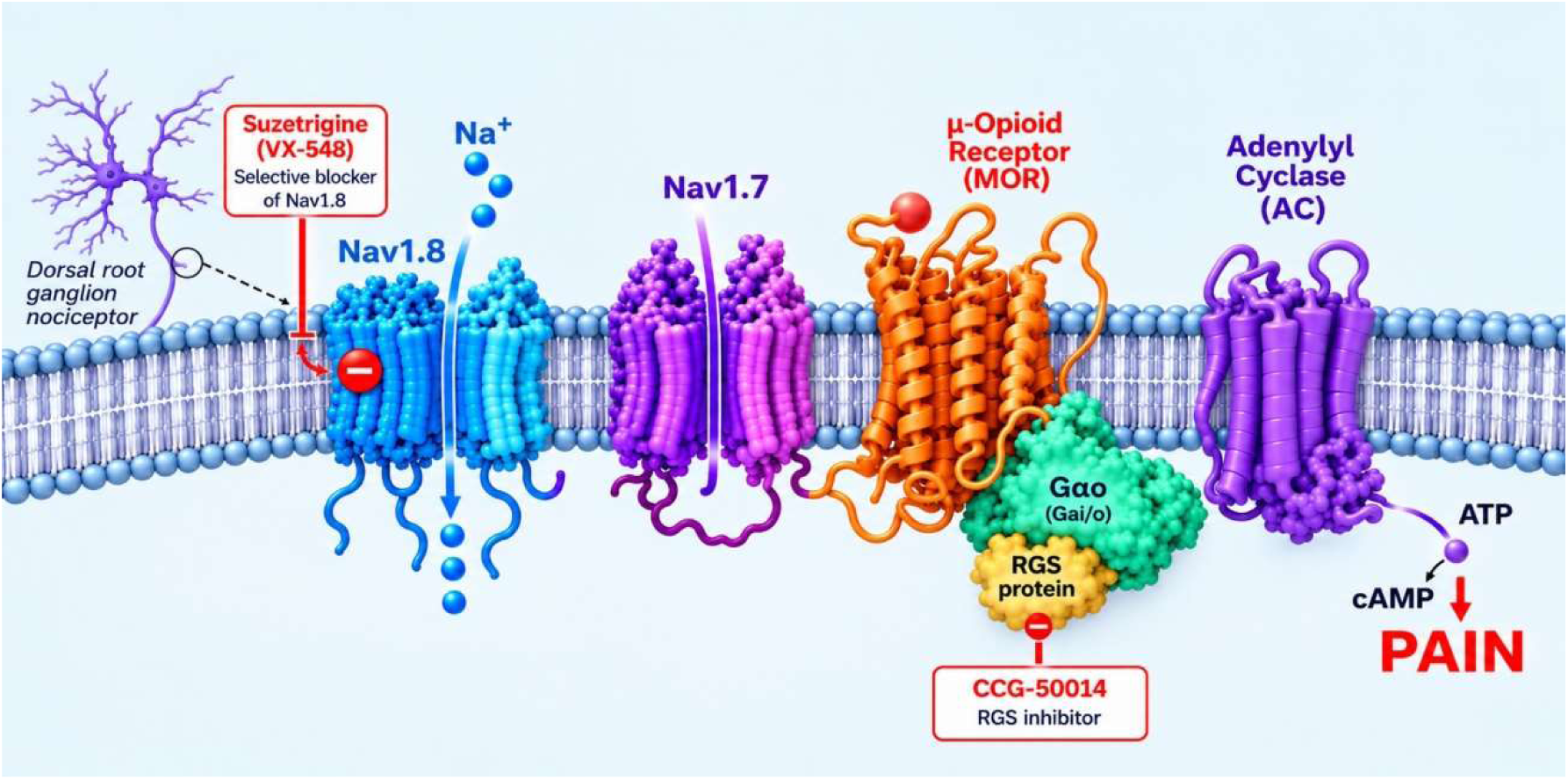
Voltage-gated sodium channels Nav1.7 and Nav1.8 and pain induction. Nav1.8 enriched in axons carries action potentials to the terminals of sensory neurons where Nav1.7 is expressed and associates with Gα0. This G-protein represses adenylyl cyclase in response to endogenous and exogenous opioids that activate opioid receptors and block pain. The endogenous GTPase activity that turns off the G-protein is activated by RGS proteins. When these are inhibited by compounds like CCG-50014, pain is lowered more effectively.

Development of an orally active congener of CCG-50014 would be very useful. The present experiments were carried out in inbred animals that may not necessarily translate to heterogenous human chronic pain conditions If they do, then suzetrigine plus 0.1mg hydrocodone should provide useful analgesia in humans at an opioid dose which lowers the potential for side effects of respiratory depression, constipation and addiction. This would be a useful opioid-sparing advance The availability of humanised Nav1.8 rats and mice that are under development will in the future provide supporting information for human trials. The present experiments nonetheless provide a rational basis for POC clinical trials to test the potential utility of combinatorial therapy with Nav1.8 blockade and Nav1.7 null opioid mechanisms that contribute to human analgesia. Other combinatorial approaches, for example NSAIDs or gabapentinoids plus suzetrigine are also worthy topics of investigation. The combination of Nav1.8 and Nav1.7 downstream opioid mechanisms as targets for analgesia is nonetheless intellectually compelling.

## Acknowledgements

Conceptualization: JNW, Funding acquisition: JNW Investigation: OD,EV,SG,AL,JI TH Project administration: JNW; Supervision: JNW TH Writing – original draft: JNW Writing – review & editing: JNW, EV OD TH JI. EV and OD are equal first authors. We thank Geoff Woods for very helpful discussions

We thank Cancer Research UK for support.

*Suzetrigine-evoked analgesia is potentiated by Naᵥ1.7 null-related mechanisms*

## Materials and Methods

### Animals

Experiments used male and female C57BL/6J mice (Charles River), 6–7 weeks old at the time of testing. Mice were housed in groups of 2–5 per cage on a 12-hour light/dark cycle with free access to water and a standard diet.

All experiments were performed with the approval of personal and project licences (PP3706054) from the United Kingdom Home Office, according to guidelines set by the Animals (Scientific Procedures) Act 1986 Amendment Regulations 2012 and the guidelines of the Committee for Research and Ethical Issues of the International Association for the Study of Pain (IASP).

### Drugs

**CCG-50014** (Apollo Scientific, cat. BUP05946) was dissolved in 12% dimethylsulfoxide (DMSO) in PBS and administered intrathecally as 100 nmol in 5 µl (20 mM stock) under isoflurane anaesthesia (1.5–3%).

**Suzetrigine** (Apollo Scientific, cat. BUP03027) was dissolved in DMSO and administered intraperitoneally at 10 mg kg^−1^ (2.5 mg ml^−1^; 0.25 mg per 25 g mouse).

**Morphine sulphate** (supplied via UCL veterinary route) was dissolved in saline and administered intraperitoneally at 0.3 mg kg^−1^ (0.0075 mg per mouse in 100 µl; 0.075 mg ml^−1^).

### Morphine dose rationale

Morphine was used at 0.3 mg kg^−1^, below the reported antinociceptive range in mice, so that any benefit of the triple combination could not be attributed to morphine alone. Doses of 0.2–2 mg kg^−1^ intraperitoneally are characterised as sub-analgesic in mice and are used in potentiation studies for that reason; 2 mg kg^−1^ i.p. produces no measurable effect in the tail-flick test when given alone; and the mouse tail-flick ED_50_ for morphine is approximately 3.2–5.6 mg kg^−1^ subcutaneously. For internal comparison, this laboratory uses 15 mg kg^−1^ s.c. as a fully analgesic reference dose in cancer-induced bone pain work.

### Dosing schedule

CCG-50014 was administered intrathecally under isoflurane anaesthesia. Animals were allowed to recover for 1 hour, after which suzetrigine and morphine were administered intraperitoneally, and behavioural testing began 30 minutes later. CCG-50014 was therefore present for approximately 90 minutes before testing. Vehicle-treated animals received saline or DMSO vehicle by the same routes and on the same schedule as drug-treated animals, including the intrathecal injection under isoflurane anaesthesia.

### Pain models

#### Formalin test

Formalin (Merck, cat. F8775; 37% formaldehyde stock diluted in saline) was injected intraplantar in a volume of 20 µl. Licking and biting of the injected paw was recorded in 5-minute bins for 60 minutes by an observer blind to treatment. Phase I was defined as 0–10 minutes, the interphase as 10–15 minutes and phase II as 15–60 minutes; the interphase was excluded from both phase totals.

#### Complete Freund’s adjuvant

Complete Freund’s adjuvant (Sigma-Aldrich/Merck, cat. F5881) was administered undiluted (neat) and injected into the left hind paw in a volume of 20 µl, in accordance with the project licence, which permits concentrations up to 100%. Behavioural testing was performed 72 hours later. Paw thickness was measured with digital callipers.

#### Behavioural assays

##### All testing was performed by an observer blind to treatment allocation. Rotarod

Animals were placed on the rotarod at 4 rpm, which was accelerated to a maximum of 40 rpm over 90 s, with a cut-off of 120 s. Latency to fall was recorded (Stirling et al., 2005).

##### Hargreaves test

Thermal thresholds were measured by the Hargreaves method (Hargreaves et al., 1988). The hind paw was exposed to a radiant heat source and the withdrawal latency recorded, with a cut-off of 30 s to prevent tissue damage.

##### von Frey test

Punctate mechanical thresholds were measured using the up–down method (Deuis et al., 2017). Mice were habituated for 2 hours in darkened enclosures with a wire mesh floor. Testing began with a 0.4 g filament applied for 3 seconds; a weaker filament was applied following a positive response and a stronger filament following a negative response, with five further applications after the first change in response. A 2 g filament was used as the cut-off. The 50% withdrawal threshold was calculated as 10^(χ + kδ)/10,000, where χ is the log₁₀ of the final filament force expressed in units of 0.1 mg, k is the tabular value for the response pattern, and δ is the mean logarithmic interval between filaments (δ = 0.175).

##### Randall–Selitto test

Deep mechanical thresholds were measured with a Randall–Selitto analgesimeter applied to the tail, as previously described (Haroun et al., 2023). The instrument records latency to withdrawal during a linear force ramp of 20 g s−1; thresholds are reported in grams (latency × 20). In the CFA experiments this site was not inflamed. Statistics

One test was pre-specified per readout. Normality was assessed with the D’Agostino– Pearson test. Within-animal baseline-versus-drug comparisons used paired t-tests with Wilcoxon signed-rank confirmation. Between-group comparisons against vehicle used two-tailed Mann–Whitney U tests, chosen for the small group sizes, with Holm correction applied across the four treatment-versus-vehicle comparisons within each assay; both raw and adjusted P values are reported. Treatment and sex were assessed by two-way ANOVA with interaction. Data are presented as mean ± SEM with individual animals shown. α = 0.05.

#### Na^+^ Imaging

Neurons were incubated with 10 μM SBFI AM (Ion Biosciences) in 0.02% Pluronic® F-127 acid (Sigma-Aldrich), diluted in extracellular solution, for 1 h at 37 °C and 5% CO_2._ Extracellular solution contained (in mM): 140 NaCl, 4 KCl, 2 CaCl_2,_ 2 MgCl_2,_ 10 HEPES, 5 Glucose, 20 Mannitol, pH 7.4 adjusted with NaOH as previously described (*1*). Prior to the experiment, coverslips were washed in extracellular solution to remove excess SBFI and then mounted in an RC-25 chamber (Warner Instruments) before being placed on an inverted Nikon Eclipse TE300 microscope. Images were acquired with OptoFluor® imaging software (Cairn Imaging Series 7.7) and Q Imaging EXi Aqua camera using alternate excitation at wavelengths of 340 and 380 nm, and 505 nm emission was collected separately to determine the 340/380 ratio.

For [Na]_i c_alibration, cells were exposed to different extracellular solutions with specific Na^+^ concentrations (0, 5, 10, 15 mM) for 30 min. Gramicidin at 5 μM (Sigma) and Monensin at 10 μM (Sigma) were added to these solutions to equilibrate extracellular and intracellular concentrations. To reduce variability between coverslips, an intracellular calibration of sodium was performed post-acquisition for each experiment with 5 and 15 mM solutions. Calibration solutions contained (in mM): 130 Na^+^ + K^+^, 30 Cl^-^, 0.6 mM MgCl_2,_ 0.5 mM CaCl_2,_ and 100 gluconate (pH 7.4).

##### Acquisition

Mice expressing GCaMP3 or GCaMP6f (8 to 14 weeks, male and female) were anesthetized using ketamine (120 mg/kg) and medetomidine (1.2 mg/kg). Depth of anaesthesia was confirmed by pedal reflex and breathing rate. Animals were maintained at a constant body temperature of 37°C using a heated mat (VetTech). Lateral laminectomy was performed at spinal level L3-5. In brief, the skin was incised longitudinally, and the paravertebral muscles were cut to expose the vertebral column. Transverse and superior articular processes of the vertebra were removed using microdissection scissors and OmniDrill 35 (WPI). To obtain a clear image of the sensory neuron cell bodies in the ipsilateral dorsal root ganglion (DRG), the dura mater and the arachnoid membranes were carefully opened using microdissection forceps. The animal was mounted onto a custom-made clamp attached to the vertebral column (L1), rostral to the laminectomy. The trunk of the animal was slightly elevated to minimize interference caused by respiration. Artificial cerebrospinal fluid [containing 120 mM NaCl, 3 mM KCl, 1.1 mM CaCl_2,_ 10 mM glucose, 0.6 mM NaH_2P_O_4,_ 0.8 mM MgSO_4,_ 18 mM NaHCO_3 (_pH 7.4) with NaOH] was perfused over the exposed DRG during the procedure to maintain tissue integrity, or the DRG was isolated by coating with silicone elastomer.

Images were acquired using a Leica SP8 confocal microscope (Leica). GCaMP3 or GCaMP6f was excited using a 488 nm laser (1-15% laser power). Images were acquired at 800Hz, with bidirectional laser scan. Typically, the pinhole was kept at 1 A.U, but in some experiments was increased to 1.5 A.U. to enhance brightness. Images magnification was between 0.75x and 3x optical zoom on a 10x air objective, depending on DRG anatomy. Noxious and innocuous stimuli were applied to the left hindpaw, ipsilateral to the exposed DRG. For thermal stimuli, the paw was immersed with ice-water (0°C) or water heated to 37°C or 55°C using a Pasteur pipette. For mechanical stimuli, we used noxious pinch with serrated forceps. PGE2 (500 µM in saline) was applied to the paw by intraplantar injection. Naloxone (2 mg/kg in saline) was delivered by subcutaneous injection into the scruff of the neck.

##### Analysis

Image stacks were registered to the first frame in the series using the FIJI plugin TurboReg (accurate rigid body transformation) to correct for XY drift. Stacks that showed excessive Z movement were excluded from analysis. Regions of interest (ROI) were manually drawn around apparently responding cells using the free hand tool in FIJI. Mean pixel intensity over time for each ROI was extracted and analysed. The time series of mean pixel intensity for each ROI was smoothened by a four time point moving average to remove high-frequency noise. Next, we calculated the derivative of the mean pixel intensity. We calculated a mean baseline derivative for the 10 s preceding stimulus application. Neurons were classed as responders if, within 30 s of stimulus application, the maximum derivative was greater than the baseline derivative plus five standard deviations – that is, a Z-score of at least 5. We then calculated the ΔF/F_0 v_alue for each response to obtain a normalized measure of change in fluorescence. Neurons which showed a ΔF/F_0 l_ess than 0.25 were then discarded. Each trace was then manually screened as a further precaution against false positives. The remaining neurons that made up the responding population were then used for statistical analysis.

### *In vivo* antisense oligonucleotide (ASO) treatment

Antisense oligodeoxynucleotides (ASOs) were dissolved in saline at a concentration of 15 μg per 5 μl). Adult C57BL/6J Na_V1_.7 KO mice (>8 weeks) were anaesthetised with 2 % isoflurane and received intrathecal injections at the lumbar level (L3–L6). Littermate mice expressing Na_V1_.7 were used as a control. ASOs were injected intrathecally into mice at days 1, 2, 3 and 5. DRG were isolated 72h after the last injection. An ASO targeting mouse Scn1a (Scn1a ASO: CTGTTTGCTCCATCTTGTCA) and a nontargeting control ASO (Scrambled ASO: GTTCTATTCGATCTCGTCTC) were used for the studies. All ASOs had a phosphorothioate backbone.

### DRG neuron cultures

Mice were sacrificed by inhalation of CO_2 o_verdose followed by cervical dislocation to confirm death. The vertebral column was then excised and DRG were then isolated with the use of a stereo microscope and placed in a Petri dish with Dulbecco’s phosphate buffered saline (DPBS, Sigma-Aldrich, St. Louis, Missouri). Afterward, the DRG were digested with collagenase type XI (% (w/v), Sigma-Aldrich) in DMEM (Gibco) with 1 % penicillin-streptomycin (P/S, 5000 U/mL, Invitrogen, Waltham, Massachussets) at 37 ⁰C for 1 h in 5 % CO_2._ Then, using a pipette the DRG were mechanically dissociated and placed into DMEM medium containing 10 % of fetal bovine serum (FBS, Invitrogen) and 1 % P/S. This cell suspension was centrifuged (5 min x300 g) to remove rests of connective tissue. The cell pellet was resuspended in fresh medium and seeded in the plates. One hour after cell seeding, DMEM medium was changed to Neurobasal-A medium (Gibco) supplemented with 1 % P/S, B-27® Supplement (2 %; Gibco) and GlutaMAX^TM^ (1 %; Invitrogen) and nerve growth factor (NGF, 5 ng·mL^−1^; Sigma-Aldrich). Cell cultures were kept at 37 ⁰C in a humidified incubator with 5 % CO_2._

#### *In vivo* calcium imaging

- **Acquisition**:

Mice expressing GCaMP3 (> 6 weeks, male and female) were anesthetized using ketamine (100 mg/kg), xylazine (15 mg/kg), and acepromazine (2.5 mg/kg). The depth of anaesthesia was confirmed by the pedal reflex and breathing rate. Animals were maintained at a constant body temperature of 37°C using a heated mat (VetTech). Lateral laminectomy was performed at spinal level L3-5. In brief, the skin was incised longitudinally, and the paravertebral muscles were cut to expose the vertebral column. Transverse and superior articular processes of the vertebra were removed using microdissection scissors and OmniDrill 35 (WPI). To obtain a clear image of the sensory neuron cell bodies in the ipsilateral dorsal root ganglion (DRG), the dura mater and the arachnoid membranes were carefully opened using microdissection forceps. The animal was mounted onto a custom-made clamp attached to the vertebral column (L1), rostral to the laminectomy. The trunk of the animal was slightly elevated to minimize interference caused by respiration. Artificial cerebrospinal fluid [containing 120 mM NaCl, 3 mM KCl, 1.1 mM CaCl_2,_ 10 mM glucose, 0.6 mM NaH_2P_O_4,_ 0.8 mM MgSO_4,_ 18 mM NaHCO_3 (_pH 7.4) with NaOH] was perfused over the exposed DRG during the procedure to maintain tissue integrity, or the DRG was isolated by coating with silicone elastomer.

Noxious and innocuous stimuli were applied to the left hind paw, ipsilateral to the exposed DRG. For thermal stimuli, the paw was immersed with ice-water (0°C) or water heated to 55°C using a Pasteur pipette. For mechanical stimuli, we used noxious pinch with serrated forceps. Each stimulus application was separated by an interval of at least 1 min.

Images were acquired using a Leica SP8 confocal microscope (Leica). GCaMP3 was excited using a 488 nm laser (1-15% laser power). Images were acquired at 800Hz, with bidirectional laser scan. The pinhole was kept at 1 A.U. Images magnification was between 0.75x and 3x optical zoom on a 10x air objective, depending on DRG anatomy.

- **Analysis**:

Image stacks were registered to the first frame in the series using the FIJI plugin TurboReg (accurate rigid body transformation) to correct for XY drift. Stacks that showed excessive Z movement were excluded from analysis. Regions of interest (ROI) were manually drawn around responding cells using the free hand tool in FIJI. Mean pixel intensity over time for each ROI was extracted and analysed. The time series of mean pixel intensity for each ROI was smoothened by a four-time point moving average to remove high-frequency noise. Next, we calculated the derivative of the mean pixel intensity. We calculated a mean baseline derivative for the 10 s preceding stimulus application. Neurons were classed as responders if, within 30 s of stimulus application, the maximum derivative was greater than the baseline derivative plus five standard deviations – that is, a Z-score of at least 5. We then calculated the ΔF/F_0 v_alue for each response to obtain a normalized measure of change in fluorescence. Neurons showing a ΔF/F_0 l_ess than 0.25 were then discarded. Each trace was then manually screened as a further precaution against false positives. The remaining neurons that made up the responding population were then used for statistical analysis.

### Patch-clamp recordings

Whole-cell patch-clamp recordings were conducted on small-diameter (<25 μm) DRG neurons seeded on 12-mm-diameter glass coverslips at room temperature. For cell culturing, glass coverslips were coated with poly-L-lysine (50 μg·mL^−1^, Sigma-Aldrich) for 2 h and washed four times with deionized water. Crystals were allowed to dry in a sterile air flow cabinet and then they were incubated during 1 h at 37 °C with laminin diluted in DMEM medium at a concentration of 10 μg·mL^−1^ (Sigma-Aldrich). Then, laminin was replaced by the cell suspension diluted in DMEM 10 % FBS 1 % P/S. Pulled borosilicate glass pipettes (2–5 MΩ) with OD 1.5 mm x ID 1.17 mm (Warner Instruments, Hamden, CT) were fabricated using a Narishige PC-10 puller (NARISHIGE Group, Japan). Seal resistance was ≥ 1 GΩ. The pipette internal solution contained (in mM): 144 KCl, 2 MgCl_2,_ 10 HEPES, 5 EGTA, pH 7.2 adjusted with KOH. The external solution contained (in mM): 140 NaCl, 4 KCl, 2 CaCl_2,_ 2 MgCl_2,_ 10 HEPES, 5 Glucose, 20 Mannitol, pH 7.4 adjusted with NaOH. After establishing whole-cell access, cells were recorded in current-clamp mode and acquired at a sampling frequency of 25 kHz. The firing frequency was evaluated with 1 s-current depolarizing pulses from 0 to 100 pA in 10 pA intervals. Voltage recordings were acquired with an Axopatch 200B amplifier controlled by Clampex 10.6 software and analyzed in Clampfit 10.6 (Molecular Devices, LLC., San Jose, CA). Only one cell per dish was recorded.

---

**Supplementary Figure 1.**
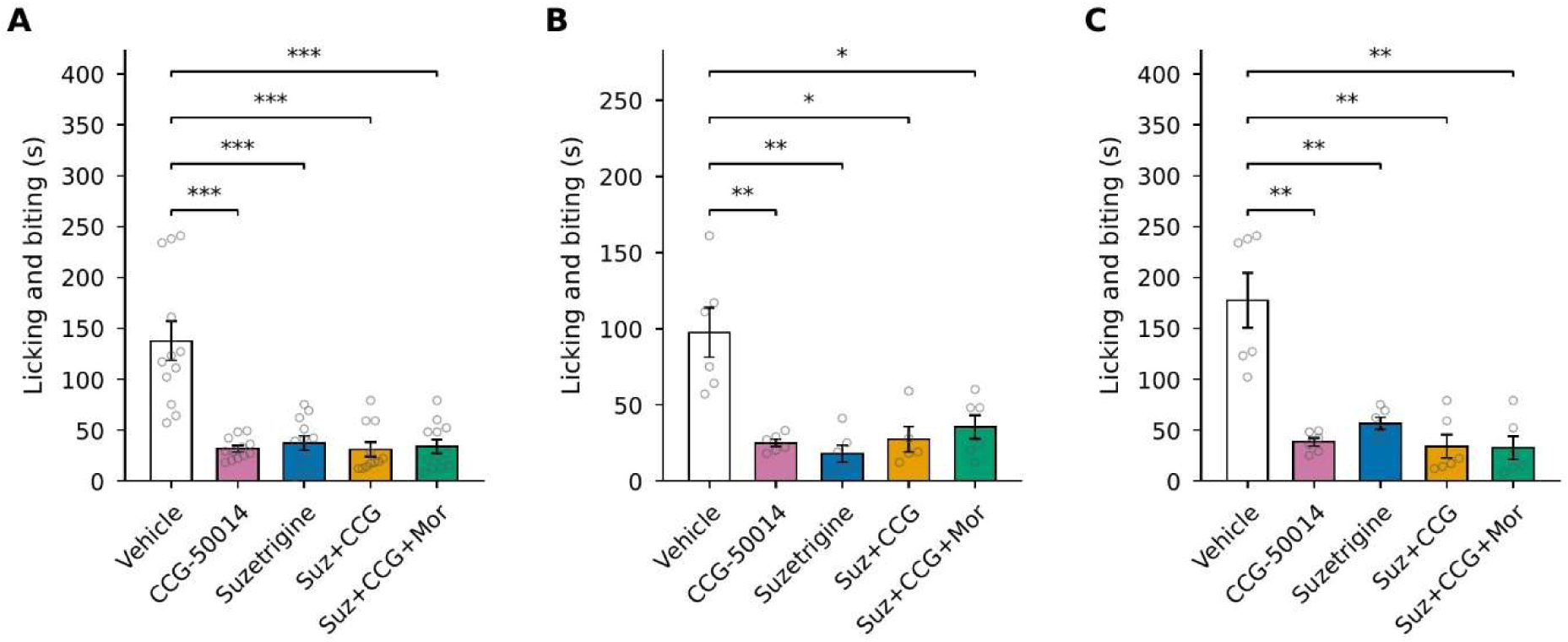
Antisense oligonucleotides knock down Scn1a (Naᵥ1.1) in sensory neurons without altering mechanical withdrawal thresholds. (A) Scn1a mRNA expression in dorsal root ganglia after intrathecal treatment with scrambled or Naᵥ1.1 antisense oligonucleotide (ASO). Bars show mean ± SD; points show individual animals. *P < 0.05. (B) Withdrawal threshold before (baseline) and after (treatment) scrambled or Naᵥ1.1 ASO, with paired measurements joined by lines.

**Supplementary Figure 2.**
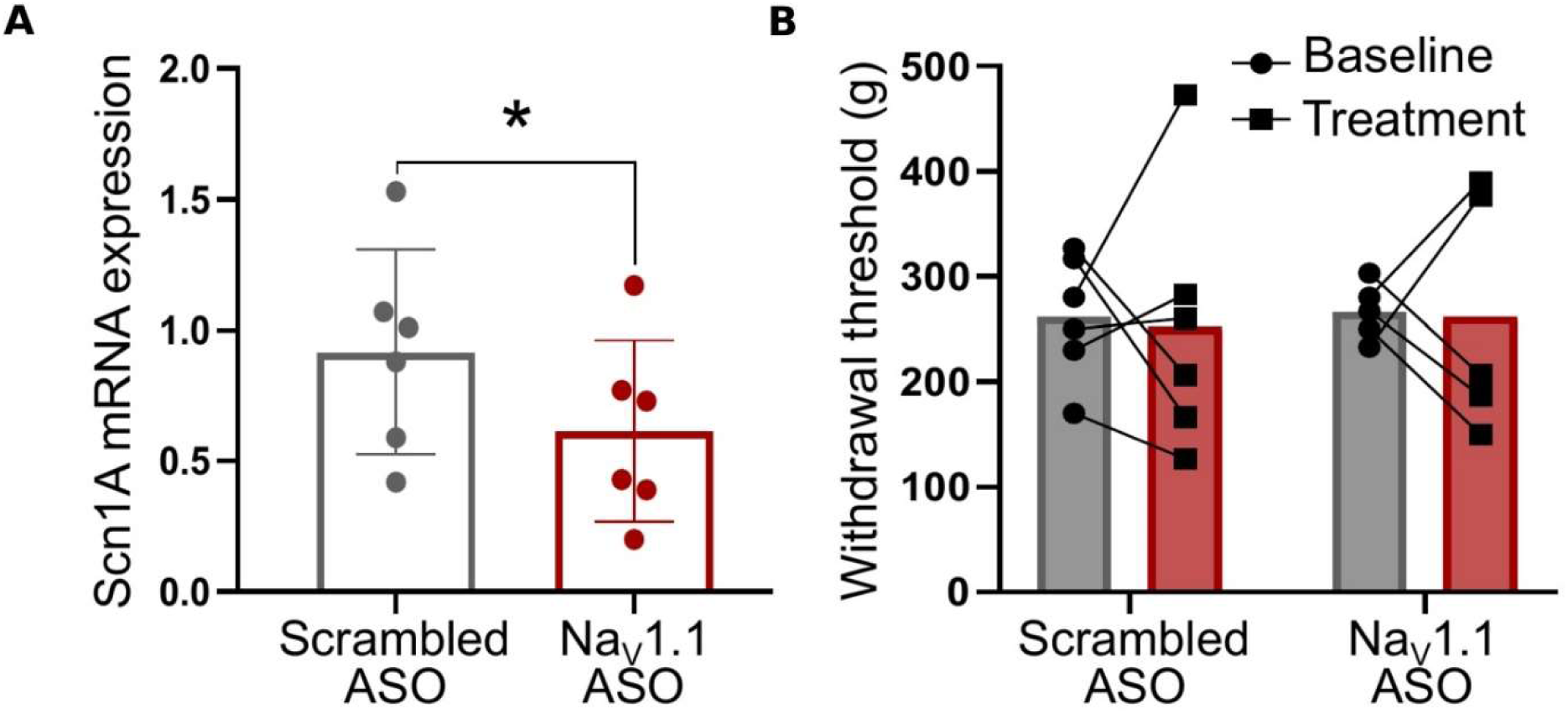
Formalin phase I responses split by sex. Total licking and biting during phase I (0 to 10 min) for (A) all animals, (B) male mice and (C) female mice. n = 5 to 6 per group per sex. Bars show the mean, error bars show the SEM and points show individual animals. Two-way ANOVA: sex P = 0.0009; treatment × sex P = 0.010. Asterisks denote two-tailed Mann–Whitney U versus vehicle within each panel, Holm-adjusted across the four comparisons. **P < 0.01; ***P < 0.001.

**Table S1.**
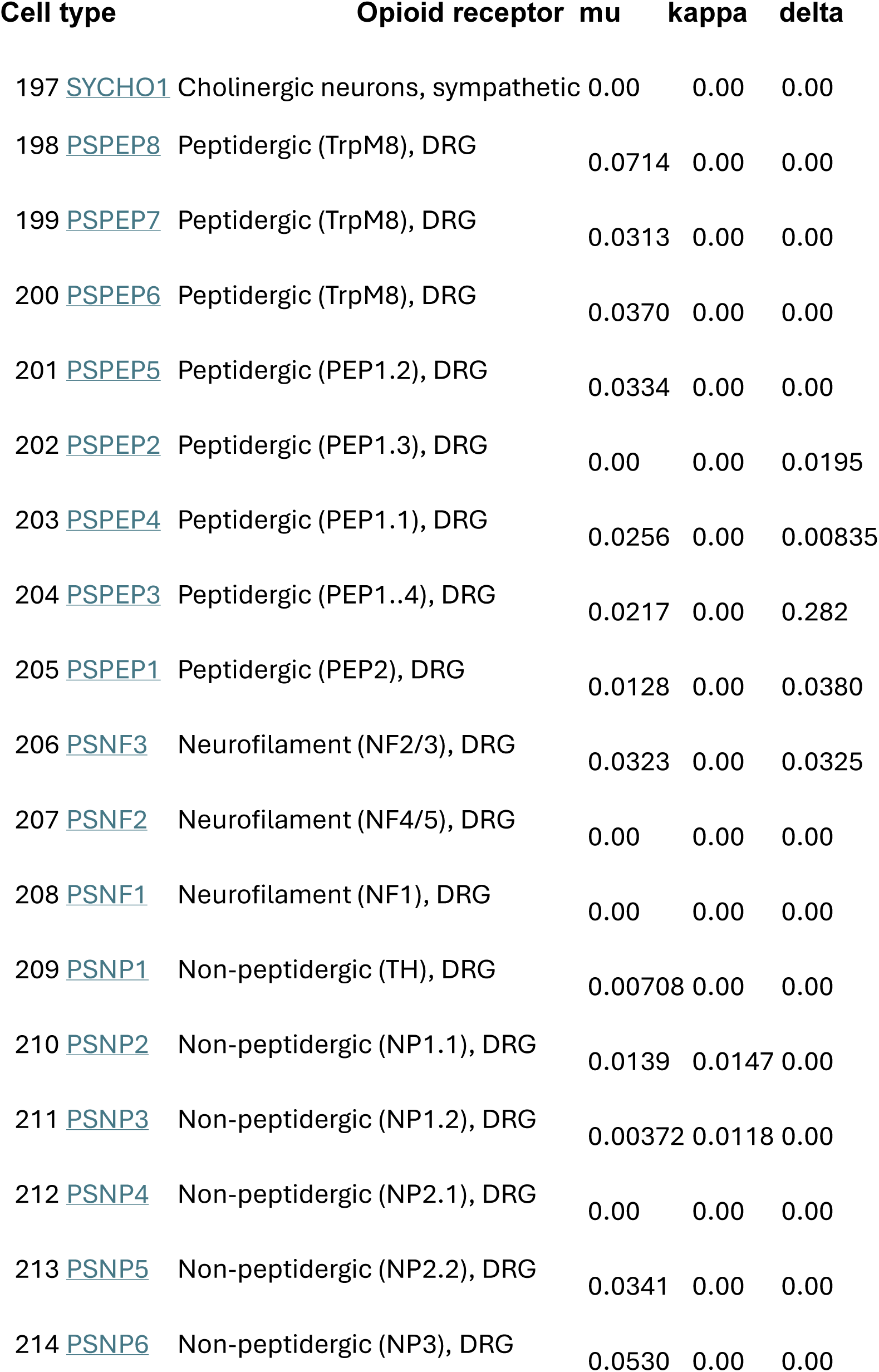
RNAseq data for sympathetic and sensory neurons shows that opioid receptors are mainly exprresed in nociceptive sensory neurons 13.

|  |  |  |  |  |
| --- | --- | --- | --- | --- |
| Cell type | Opioid receptor | mu | kappa | delta |

|  |  |  |  |  |  |
| --- | --- | --- | --- | --- | --- |
| 197 | SYCHO1 | Cholinergic neurons, sympathetic | 0.00 | 0.00 | 0.00 |
| 198 | PSPEP8 | Peptidergic (TrpM8), DRG | 0.0714 | 0.00 | 0.00 |
| 199 | PSPEP7 | Peptidergic (TrpM8), DRG | 0.0313 | 0.00 | 0.00 |
| 200 | PSPEP6 | Peptidergic (TrpM8), DRG | 0.0370 | 0.00 | 0.00 |
| 201 | PSPEP5 | Peptidergic (PEP1.2), DRG | 0.0334 | 0.00 | 0.00 |
| 202 | PSPEP2 | Peptidergic (PEP1.3), DRG | 0.00 | 0.00 | 0.0195 |
| 203 | PSPEP4 | Peptidergic (PEP1.1), DRG | 0.0256 | 0.00 | 0.00835 |
| 204 | PSPEP3 | Peptidergic (PEP1..4), DRG | 0.0217 | 0.00 | 0.282 |
| 205 | PSPEP1 | Peptidergic (PEP2), DRG | 0.0128 | 0.00 | 0.0380 |
| 206 | PSNF3 | Neurofilament (NF2/3), DRG | 0.0323 | 0.00 | 0.0325 |
| 207 | PSNF2 | Neurofilament (NF4/5), DRG | 0.00 | 0.00 | 0.00 |
| 208 | PSNF1 | Neurofilament (NF1), DRG | 0.00 | 0.00 | 0.00 |
| 209 | PSNP1 | Non-peptidergic (TH), DRG | 0.00708 | 0.00 | 0.00 |
| 210 | PSNP2 | Non-peptidergic (NP1.1), DRG | 0.0139 | 0.0147 | 0.00 |
| 211 | PSNP3 | Non-peptidergic (NP1.2), DRG | 0.00372 | 0.0118 | 0.00 |
| 212 | PSNP4 | Non-peptidergic (NP2.1), DRG | 0.00 | 0.00 | 0.00 |
| 213 | PSNP5 | Non-peptidergic (NP2.2), DRG | 0.0341 | 0.00 | 0.00 |
| 214 | PSNP6 | Non-peptidergic (NP3), DRG | 0.0530 | 0.00 | 0.00 |

## References

1. Wood, J.N. (2021). The Oxford Handbook of The Neurobiology of Pain (OUP).

2. Gordh, T. (2010). Lidocaine: the origin of a modern local anesthetic. 1949. Anesthesiology 113, 1433–1437. 10.1097/ALN.0b013e3181fcef48.

3. Taylor, C.P., and Harris, E.W. (2020). Analgesia with Gabapentin and Pregabalin May Involve N-Methyl-d-Aspartate Receptors, Neurexins, and Thrombospondins. J Pharmacol Exp Ther 374, 161–174. 10.1124/jpet.120.266056.

4. Judd, D., King, C.R., and Galke, C. (2023). The Opioid Epidemic: A Review of the Contributing Factors, Negative Consequences, and Best Practices. Cureus 15, e41621. 10.7759/cureus.41621.

5. Osteen, J.D., Immani, S., Tapley, T.L., Indersmitten, T., Hurst, N.W., Healey, T., Aertgeerts, K., Negulescu, P.A., and Lechner, S.M. (2025). Pharmacology and Mechanism of Action of Suzetrigine, a Potent and Selective Na(V)1.8 Pain Signal Inhibitor for the Treatment of Moderate to Severe Pain. Pain Ther 14, 655–674. 10.1007/s40122-024-00697-0.

6. Hasoon, J., Nguyen, A., Govindaraj, R., and Robinson, C.L. (2026). Suzetrigine for Cancer-Related Bone Pain: A Three-Patient Case Series. Orthop Rev (Pavia) 18, 162158. 10.52965/001c.162158.

7. Ali, M.Y., Antunes, F.T.T., Huang, S., Chen, L., and Zamponi, G.W. (2025). Pharmacological inhibition of NaV1.8 by suzetrigine reveals potent analgesic potential without tolerance development in mice. Mol Brain 18, 86. 10.1186/s13041-025-01253-3.

8. Cox, J.J., Reimann, F., Nicholas, A.K., Thornton, G., Roberts, E., Springell, K., Karbani, G., Jafri, H., Mannan, J., Raashid, Y., et al. (2006). An SCN9A channelopathy causes congenital inability to experience pain. Nature 444, 894–898. nature05413 [pii] 10.1038/nature05413.

9. MacDonald, D.I., Sikandar, S., Weiss, J., Pyrski, M., Luiz, A.P., Millet, Q., Emery, E.C., Mancini, F., Iannetti, G.D., Alles, S.R.A., et al. (2021). A central mechanism of analgesia in mice and humans lacking the sodium channel Na(V)1.7. Neuron 109, 1497–1512 e1496. 10.1016/j.neuron.2021.03.012.

10. Regan, C.P., Morissette, P., Kraus, R.L., Wang, E., Arrington, L., Vavrek, M., de Hoon, J., Depre, M., Lodeweyck, T., Demeyer, I., et al. (2024). Autonomic Dysfunction Linked to Inhibition of the Na(v)1.7 Sodium Channel. Circulation 149, 1394–1396. 10.1161/CIRCULATIONAHA.123.067331.

11. Zhou, X., Zhou, X., Ji, M., Xu, P., Cox, J.J., Zhao, J., and Zhang, C. (2026). Intrathecal administration of the Nav1.7 inhibitor PF-05089771 produces rapid and side-effect-free analgesia in mice. Pain 167, 589–605. 10.1097/j.pain.0000000000003823.

12. Weiss, J., Pyrski, M., Jacobi, E., Bufe, B., Willnecker, V., Schick, B., Zizzari, P., Gossage, S.J., Greer, C.A., Leinders-Zufall, T., et al. (2011). Loss-of-function mutations in sodium channel Nav1.7 cause anosmia. Nature 472, 186–190. 10.1038/nature09975.

13. Usoskin, D., Furlan, A., Islam, S., Abdo, H., Lonnerberg, P., Lou, D., Hjerling-Leffler, J., Haeggstrom, J., Kharchenko, O., Kharchenko, P.V., et al. (2015). Unbiased classification of sensory neuron types by large-scale single-cell RNA sequencing. Nat Neurosci 18, 145–153. 10.1038/nn.3881.

14. Clark, M.J., Harrison, C., Zhong, H., Neubig, R.R., and Traynor, J.R. (2003). Endogenous RGS protein action modulates mu-opioid signaling through Galphao. Effects on adenylyl cyclase, extracellular signal-regulated kinases, and intracellular calcium pathways. J Biol Chem 278, 9418–9425. 10.1074/jbc.M208885200.

15. Lopez-Mateos, D., Harris, B.J., Hernandez-Gonzalez, A., Yarov-Yarovoy, V., and Wulff, H. (2025). Recent advances in the pharmacology of voltage-gated ion channels. Pharmacol Rev 77, 100090. 10.1016/j.pharmr.2025.100090.

16. Iseppon, F., Kanellopoulos, A.H., Tian, N., Zhou, J., Caan, G., Chiozzi, R., Thalassinos, K., Cubuk, C., Lewis, M.J., Cox, J.J., et al. (2024). Sodium channels Na(v)1.7, Na(v)1.8 and pain; two distinct mechanisms for Na(v)1.7 null analgesia. Neurobiol Pain 16, 100168. 10.1016/j.ynpai.2024.100168.

17. Wood, J.N., Yan, N., Huang, J., Zhao, J., Akopian, A., Cox, J.J., Woods, C.G., and Nassar, M.A. (2025). Sensory neuron sodium channels as pain targets; from cocaine to Journavx (VX-548, suzetrigine). J Gen Physiol 157. 10.1085/jgp.202513778.

18. Pereira, V., Millet, Q., Aramburu, J., Lopez-Rodriguez, C., Gaveriaux-Ruff, C., and Wood, J.N. (2018). Analgesia linked to Nav1.7 loss of function requires micro- and delta-opioid receptors. Wellcome Open Res 3, 101. 10.12688/wellcomeopenres.14687.1.

19. Minett, M.S., Pereira, V., Sikandar, S., Matsuyama, A., Lolignier, S., Kanellopoulos, A.H., Mancini, F., Iannetti, G.D., Bogdanov, Y.D., Santana-Varela, S., et al. (2015). Endogenous opioids contribute to insensitivity to pain in humans and mice lacking sodium channel Nav1.7. Nat Commun 6, 8967-8967, 8967. 10.1038/ncomms9967.

20. Isensee, J., Krahe, L., Moeller, K., Pereira, V., Sexton, J.E., Sun, X., Emery, E., Wood, J.N., and Hucho, T. (2017). Synergistic regulation of serotonin and opioid signaling contributes to pain insensitivity in Nav1.7 knockout mice. Sci Signal 10. 10.1126/scisignal.aah4874.

21. Cai, S., Moutal, A., Yu, J., Chew, L.A., Isensee, J., Chawla, R., Gomez, K., Luo, S., Zhou, Y., Chefdeville, A., et al. (2021). Selective targeting of NaV1.7 via inhibition of the CRMP2-Ubc9 interaction reduces pain in rodents. Sci Transl Med 13, eabh1314. 10.1126/scitranslmed.abh1314.

22. Senese, N.B., Kandasamy, R., Kochan, K.E., and Traynor, J.R. (2020). Regulator of G-Protein Signaling (RGS) Protein Modulation of Opioid Receptor Signaling as a Potential Target for Pain Management. Front Mol Neurosci 13, 5. 10.3389/fnmol.2020.00005.

23. Avrampou, K., Pryce, K.D., Ramakrishnan, A., Sakloth, F., Gaspari, S., Serafini, R.A., Mitsi, V., Polizu, C., Swartz, C., Ligas, B., et al. (2019). RGS4 Maintains Chronic Pain Symptoms in Rodent Models. J Neurosci 39, 8291–8304. 10.1523/JNEUROSCI.3154-18.2019.

24. Minett, M.S., Pereira, V., Sikandar, S., Matsuyama, A., Lolignier, S., Konstantinos, A., Mancini, F., Iannetti, G.D., Bogdanov, Y.D., Santana-Varela, S., et al. (2015). Endogenous opioids contribute to insensitivity to pain in humans and mice lacking sodium channel Nav1.7. Nature Communications.

25. Yoon, S.Y., Woo, J., Park, J.O., Choi, E.J., Shin, H.S., Roh, D.H., and Kim, K.S. (2015). Intrathecal RGS4 inhibitor, CCG50014, reduces nociceptive responses and enhances opioid-mediated analgesic effects in the mouse formalin test. Anesth Analg 120, 671–677. 10.1213/ANE.0000000000000607.

26. Gear, R.W., Gordon, N.C., Hossaini-Zadeh, M., Lee, J.S., Miaskowski, C., Paul, S.M., and Levine, J.D. (2008). A subanalgesic dose of morphine eliminates nalbuphine anti-analgesia in postoperative pain. J Pain 9, 337–341. 10.1016/j.jpain.2007.11.011.

27. Harris, H.M., Gul, W., ElSohly, M.A., and Sufka, K.J. (2018). Effects of Cannabidiol and a Novel Cannabidiol Analog against Tactile Allodynia in a Murine Model of Cisplatin-Induced Neuropathy: Enhanced Effects of Sub-Analgesic Doses of Morphine. Med Cannabis Cannabinoids 1, 54–59. 10.1159/000489077.

28. Afify, E.A., and Andijani, N.M. (2017). Potentiation of Morphine-Induced Antinociception by Propranolol: The Involvement of Dopamine and GABA Systems. Front Pharmacol 8, 794. 10.3389/fphar.2017.00794.

29. Akopian, A.N., Souslova, V., England, S., Okuse, K., Ogata, N., Ure, J., Smith, A., Kerr, B.J., McMahon, S.B., Boyce, S., et al. (1999). The tetrodotoxin-resistant sodium channel SNS has a specialized function in pain pathways. Nat Neurosci 2, 541–548. 10.1038/9195.

30. Rajasingham, R., and Qi, Y. (2025). Suzetrigine, a Non-Opioid Small-Molecule Analgesic: Mechanism of Action, Clinical, and Translational Science. Clin Transl Sci 18, e70414. 10.1111/cts.70414.

31. Fuggle, N., Laslop, A., Rizzoli, R., Al-Daghri, N., Alokail, M., Balkowiec-Iskra, E., Beaudart, C., Bruyere, O., Bemden, A.B., Burlet, N., et al. (2025). Treatment of Osteoporosis and Osteoarthritis in the Oldest Old. Drugs 85, 343–360. 10.1007/s40265-024-02138-w.

32. Branco, T., Tozer, A., Magnus, C.J., Sugino, K., Tanaka, S., Lee, A.K., Wood, J.N., and Sternson, S.M. (2016). Near-Perfect Synaptic Integration by Nav1.7 in Hypothalamic Neurons Regulates Body Weight. Cell 165, 1749–1761. 10.1016/j.cell.2016.05.019.

33. Muller, P., Draguhn, A., and Egorov, A.V. (2024). Persistent sodium currents in neurons: potential mechanisms and pharmacological blockers. Pflugers Arch 476, 1445–1473. 10.1007/s00424-024-02980-7.

34. Clatot, J., Hoshi, M., Wan, X., Liu, H., Jain, A., Shinlapawittayatorn, K., Marionneau, C., Ficker, E., Ha, T., and Deschenes, I. (2017). Voltage-gated sodium channels assemble and gate as dimers. Nat Commun 8, 2077. 10.1038/s41467-017-02262-0.

35. Jaffe, S. (2023). Sackler money to go towards reducing overdose deaths. Lancet 401, 1918. 10.1016/S0140-6736(23)01178-9.

36. Wirth, C., Mustafa, J., Terlecky, S.R., and Pilkington, B. (2026). Has the Pendulum Swung Too Far? Ethical Issues in the Conservative Use of Prescription Opioids. HEC Forum. 10.1007/s10730-026-09607-7.

